# High-Throughput Controlled Mechanical Stimulation and Functional Imaging *In Vivo*

**DOI:** 10.1101/107318

**Authors:** Yongmin Cho, Daniel A. Porto, Hyundoo Hwang, Laura J. Grundy, William R. Schafer, Hang Lu

## Abstract

Understanding mechanosensation and other sensory behavior in genetic model systems such as *C. elegans* is relevant to many human diseases. These studies conventionally require immobilization by glue and manual delivery of stimuli, leading to low experimental throughput and high variability. Here we present a microfluidic platform that delivers precise mechanical stimuli robustly. The system can be easily used in conjunction with functional imaging and optical interrogation techniques, as well as other capabilities such as sorting or more sophisticated fluid delivery schemes. The platform is fully automated, thereby greatly enhancing the throughput and robustness of experiments. We show that behavior of the well-known gentle and harsh touch neurons and their receptive fields can be recapitulated in our system. Using calcium dynamics as a readout, we demonstrate the ability to perform a drug screen *in vivo*. Furthermore, using an integrated chip platform that can deliver both mechanical and chemical stimuli, we examine sensory integration in interneurons in response to multimodal sensory inputs. We envision that this system will be able to greatly accelerate the discovery of genes and molecules involved in mechanosensation and multimodal sensory behavior, as well as the discovery of therapeutics for related diseases.

## Introduction

Mechanosensation is required for multiple sensory modalities such as touch, hearing, and balance, and is linked to a multitude of disorders including deafness^1–4^. Molecular mechanisms for mechanotransduction have been partially elucidated using a variety of model organisms, including *Caenorhabditis elegans* ^5–16^. Conventional mechanosensation experiments with *C. elegans* typically involve the manual delivery of a mechanical stimulus to anterior or posterior regions of animals via an eyebrow hair or metal pick^5,17–19^, and visual scoring of touch avoidance behavior, an assay subject to considerable variability between experimenters. Computer-controlled stimulation methods, for example using a piezo-driven micro stylus, have been used with electrophysiological and functional imaging approaches to deliver more repeatable mechanical stimuli to animals^20,21^. However, recording of neuronal responses by patch clamping or calcium imaging in response to precisely controlled mechanical stimulation requires animals to be immobilized with glue^15,20,21^, limiting experimental throughput and disallowing the recovery of animals for screens or further experimentation. Moreover, gluing itself is likely to affect neuronal or circuit response, and differences in the extent of gluing introduce additional experimental variability.

Microfluidics has long been used as a “lab-on-a-chip” technology, allowing for well-controlled and high-throughput experiments with small samples^22^. In addition to enabling precise perturbations on the micron scale, microfluidic devices can easily be designed to work together with optical microscopy, allowing for imaging of fluorescent probes such as calcium indicators. For *C. elegans* experimentation particularly, microfluidics has been a widely adopted technology due to the match in length scale and compatibility with fluid handling^23^. Various devices have been developed for delivering a variety of stimuli, including chemical cues and temperature gradients, while simultaneously recording neuronal responses through calcium imaging^24–33^. In contrast, there are currently no microfluidic devices capable of delivering mechanical stimuli to *C. elegans*, or do so while recording neuronal activities in a controlled manner. In this work we present a microfluidic platform for delivering robust and precise mechanical stimuli to *C. elegans* by using pneumatically actuated structures. The device is fully automated, minimizing human variability and improving experimental throughput; it is fully compatible with fluorescent imaging of calcium dynamics of neurons, which enables mechanistic interrogations as well as high-throughput genetic or drug screens. Furthermore, the mechanical stimulus module of the device can be easily integrated with other microfluidic modalities, allowing for multimodal stimulation for sensory integration studies. Here we demonstrate the design and utility of such a system in the context of high-throughput screening, as well as interrogate circuit dynamics in multimodal sensory behavior.

## Results

Our microfluidic device is optimized to deliver precise and repeatable mechanical stimuli to different anatomical regions of *C. elegans* (Fig. 1). After animals are loaded into an imaging channel (where the animals are not immobilized but their movement is much reduced from freely moving behavior), mechanical stimuli are delivered through two pairs of in-plane PDMS membrane structures (Fig. 1a and Supplementary Fig. 1). The structures are pressure-actuated, and when deflected, exert a mechanical stimulus on animals trapped in the imaging channel. The deflection and deformation caused by these actuations are in similar ranges as conventional approaches (Supplementary Fig. 2)^20,21^. Two additional actuated structures act as loading and imaging valves (Fig. 1a). This design retains animals in plane and relatively stationary but not fully immobilized, thus allowing high-quality imaging of calcium transients in cell bodies and subcellular processes (Fig. 1b). To automatically identify the fluorescently labeled neuron of interest and extract quantitative calcium transients, we developed a neuron tracking algorithm (Supplementary Fig. 3). The actuated structures are connected to a pressure source via individually controlled off-chip solenoid valves, allowing for an automated and rapid “load-and-image” routine (Fig. 1c). Additionally, the duration and pressure of stimuli can be easily controlled, allowing for the study of a variety of behaviors upon mechanical stimuli such as graded response, habituation, and arousal. Furthermore, this design can be easily adapted to allow for sorting and imaging animals of various sizes.

**Figure 1:**
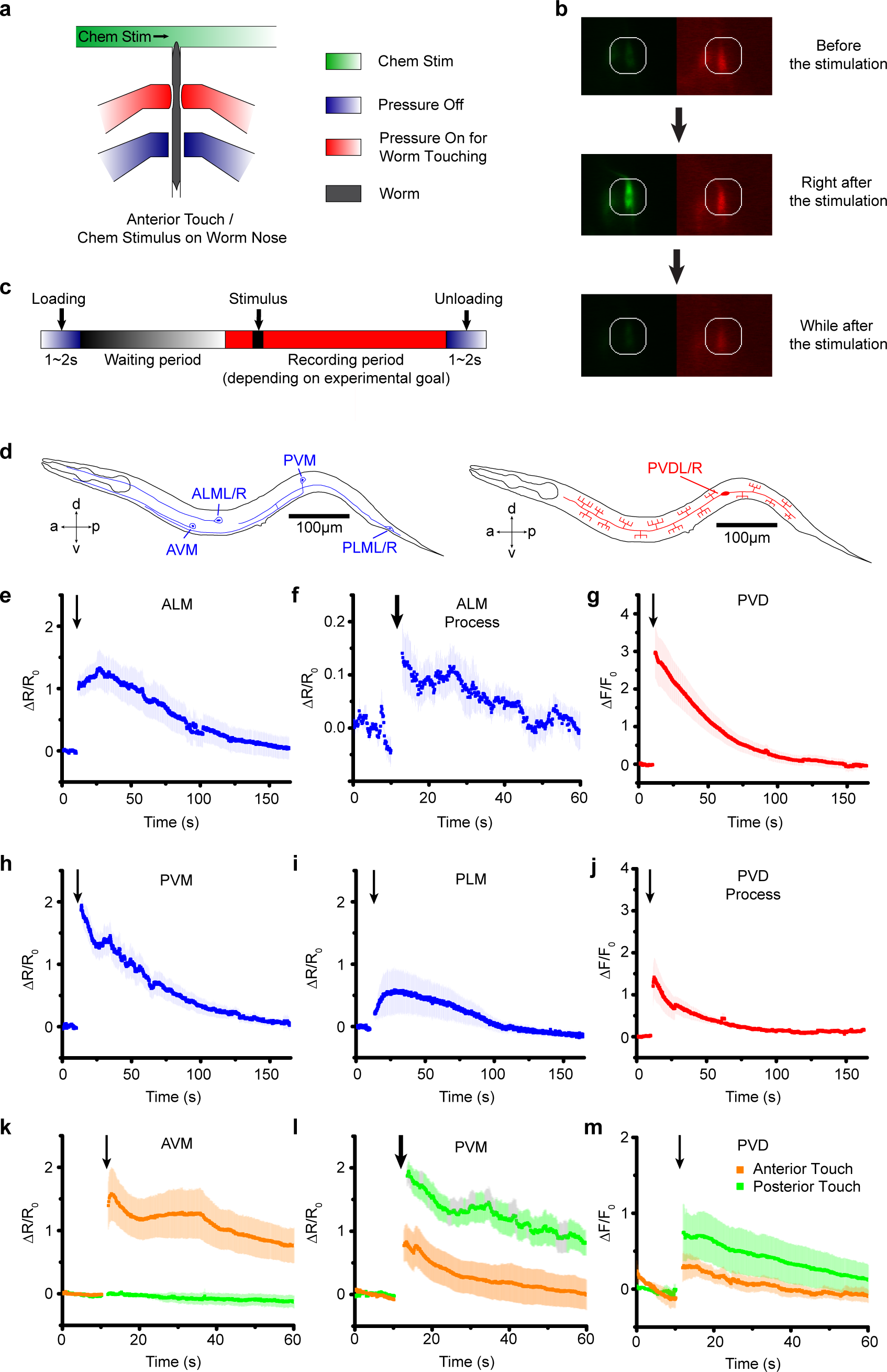
Microfluidic platform can robustly deliver mechanical stimulus and allow imaging of calcium responses in *C. elegans* mechanoreceptor neurons. **a)** The device employs multiple sets of actuated structures: valves to trap animals in a reproducible position, and two sets of actuation valves used to deliver mechanical stimuli to the anterior and posterior regions of the body. **b)** Sample frames from an activated neuron show changes in fluorescence due to mechanical stimulus. Because animals are not fully immobilized, and neurons of interest move during recordings due to the mechanical stimulus and behavioral responses, a tracking algorithm (Supplementary Fig. 3) was developed in order to automatically record the GCaMP and RFP intensities of traces from individual trials. **c)** Timeline of on-chip mechano-stimulation and functional imaging of neurons. The loading and unloading of each worm requires only a few seconds. Each animal is given a waiting period to acclimate to the environment before being stimulated and imaged. Each trial is performed by recording video to track neuronal dynamic responses and applying mechanical stimuli. **d)** Schematics of the mechanoreceptor neurons in this study: six gentle touch neurons -AVM, ALML/R, PVM, and PLML/R - and harsh touch PVDL/R neurons. **e-j)** Responses of the *C. elegans* gentle touch and harsh touch neurons to mechanical stimuli. Average traces of GCaMP6 signals in **e)** ALM soma to 1s stimulus (n=16), **f)** ALM process to 2s stimulus (n=7), **g)** PVD soma (n=9, 55psi), **h)** PVM soma (n=17), **i)** PLM soma (n=9), and **j)** PVD process to 1s stimulus (n=5) at 45psi. Error bars represent SEM. **k - m)** Gentle and harsh touch neurons exhibit reliable calcium responses when spatially resolved stimulus was delivered to the appropriate regions of animals in our system. **k)** The activity of AVM responses to 1s anterior but not posterior stimuli (anterior: n=10, posterior: n=5) at 45psi. **l, m**) Gentle touch neuron, PVM, (2s stimulation, anterior: n=10, posterior: n=11) and harsh touch neuron, PVD, (2s stimulation, anterior: n=9, posterior: n=3) respond to both anterior and posterior stimuli at 45psi. Both neurons show the higher peak of neuronal responses to posterior stimuli than anterior stimuli. Error bars represent SEM. Orange denotes anterior touch and green denotes posterior touch. For panels (e-m), arrow thickness indicates stimulation duration. 1s and 2s stimulations are represented by thin and thick arrows, respectively.

To demonstrate the utility of the system, we examined the responses of the classic gentle (AVM, ALMR/L, PVM, and PLMR/L) and harsh (PVD) touch receptor neurons^19^ (Fig. 1d). The stimulus is traditionally delivered to moving worms by a metal pick^19^, or to immobilized worms by a stiff probe^15,20^. By imaging calcium transients in animals expressing the genetically encoded calcium indicator (GECI) GCaMP6m^34^ in these touch receptor neurons, we show that the same device can deliver stimuli capable of exciting both the gentle and harsh touch neurons. Upon delivery of either a 1-or 2-second stimulus, calcium levels in cell bodies as well as in neuronal processes of both the gentle and harsh touch neurons rose as expected (Fig. 1e-m and Supplementary Movie 1-4). Spatially, these responses were consistent with the individual neurons' receptive fields as defined by anatomical and/or calcium imaging data^5,18,20,35^. AVM responded to anterior but not posterior stimuli (Fig. 1k). In contrast, PVD responded to both anterior and posterior stimuli, as did PVM, with the responses to posterior stimuli being stronger for both of these classes of neurons (Fig. 1l, m). These results demonstrate that the mechanosensory chip delivers biologically relevant, spatially well-defined stimuli.

Because our system delivers mechanical stimuli by applying externally controlled pressure to actuated structures, the stimuli can be regulated by the magnitude and duration of the applied pressure (Fig. 2 and Supplementary Fig. 2). In the range of stimuli of relevance, the deformation in the animal tissue is roughly linear to the actuation pressure (Supplementary Fig. 2). To examine the effects of these two parameters, we applied anterior stimuli of varying levels of pressure and durations, and measured calcium activity in AVM neurons (Fig. 2a-d and Supplementary Fig. 4a, 5). Peak calcium transients were roughly proportional to the pressure applied (Fig. 2a, c) and the stimulus duration (Fig. 2b, d and Supplementary Fig. 4, 5). We also tested AVM’s responses in the well-known *mec-4*/DEG/ENaC channel mutant (Fig. 2e). As expected, the *mec-4* mutant gives negligible response and is insensitive to the magnitude of the stimulation input in the gentle touch regime, but is responsive to harsh touch, perhaps even more so than wild-type (Fig. 2e,f). Similarly, in the harsh touch regime, we presented posterior stimuli of varying pressure and durations, and observed responses in PVD neurons (Fig. 2g, h). As expected, compared to gentle touch neurons, PVD required higher pressure (55 psi) or longer duration of stimulus (5s) at low pressure to elicit similar responses. Furthermore, PVD also shows graded response to pressure and duration (Fig. 2g, h and Supplementary Fig. 4b).

**Figure 2:**
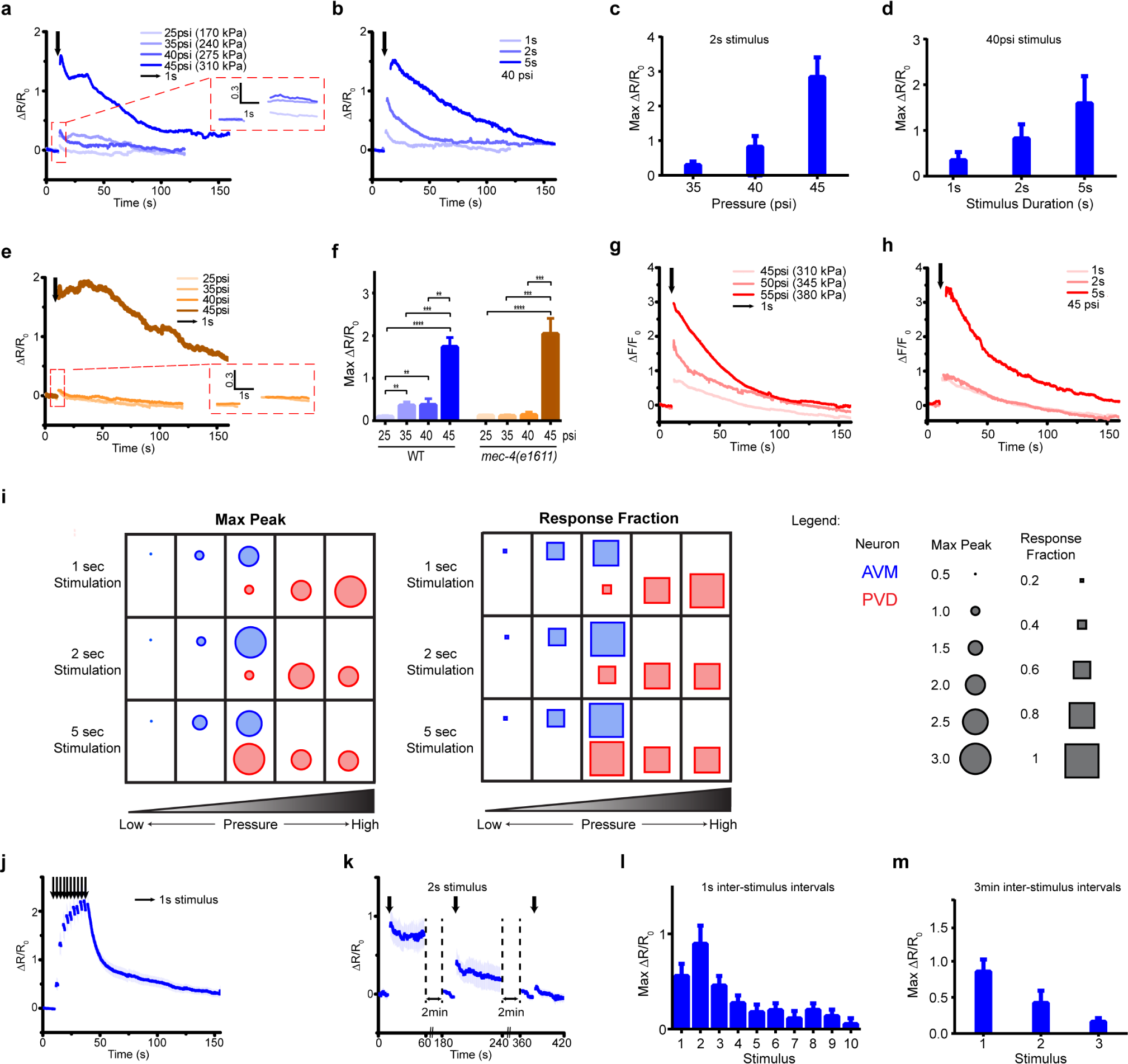
The microfluidic platform delivers mechanical stimuli emulating both gentle and harsh touch by varying the magnitude of applied pressure and duration of the stimuli. **a - b)** Average traces of GCaMP6 signal in AVM neuron in response to diverse pressures and stimulus durations. **a)** Applied 1s stimulation with diverse pressures (25 psi: n=11, 35 psi: n=25, 40 psi: n=8, 45 psi: n=27). **b)** Applied 40 psi stimulation with diverse stimulus durations (1s: n=8, 2s: n=10, 5s: n=10). **c - d)** Maximum responses of calcium transients correlate with **c)** the applied pressure (2s stimulus, 35 to 45 psi) and **d)** the duration of stimuli (1 to 5s stimuli, 40 psi). Error bars represent SEM. **e)** Average calcium responses of *mec-4(e1611)* mutants in AVM neuron to diverse pressures with 1s stimulus (25 psi: n=18, 35 psi: n=10, 40 psi: n=9, 45 psi: n=10). **f)** Maximum responses of calcium responses of wild-type and *mec-4* mutant animals (Mann-Whitney Test, * p<0.05, ** p<0.01, *** p<0.001, **** p<0.0001). **g - h)** Average traces of GCaMP6 signal in PVD neuron in response to diverse pressures and stimulus durations. **g)** Applied 1s stimulation with diverse pressures (45 psi: n=9, 50 psi: n=6, 55 psi: n=9). **h)** Applied 45 psi stimulation with diverse stimulus durations (1s: n=9, 2s: n=4, 5s: n=6). **i)** Quantitative responses of AVM and PVD under different stimulation conditions. Both gentle touch AVM neurons and harsh touch sensing PVD neurons can be stimulated using this platform when using the right parameter regime. Each column refers to the applied pressure magnitude and each row refers to the applied durations of stimulation. For each data point, the circle size indicates the max response value from 0 to 3.0 and the rectangle size indicates response fraction from 0 to 1. Response fraction is defined as the percentage of traces that show a max response value of higher than 0.5. **j-m)** Delivery of precisely repeated stimuli in AVM. **j, l)** When worms are exposed to 1s stimuli with short inter-stimulus intervals (1s), the neurons exhibited an incremental increase in response magnitude up to the second stimulus, and a reduced response in later stimuli (n=19). **k, m)** In contrast, when exposed to 2s stimuli with long inter-stimulus intervals (3 min), the response magnitude was reduced after each stimulus (n=10). Error bars represent SEM.

Interestingly, in addition to response magnitude, the response rates of both the gentle touch and the harsh touch neurons are also functions of the stimulation pressure and duration (Fig. 2i). For AVM, stimuli with actuation pressure higher than 40 psi produce a response rate (fraction of animals responding) of >90%, while below 30 psi the response is more stochastic (<20%) (Fig. 2i and Supplementary Fig. 4-6). Applying stimuli at lower actuation pressure also elicits a less sustained response or small magnitude of response and shorter stimuli elicits less response.

Besides simple stimulation, our system can also be used to deliver repeated stimuli in order to examine phenomena such as habituation and desensitization. Previous work has shown that presenting repeated mechanical stimuli can cause habituation in mechanosensory neurons^20^. To ask whether this phenomenon can be recapitulated in our system, we delivered repeated stimuli to animals using either short (1s) or long (3 min) inter-stimulus intervals (Fig. 2j-m and Supplementary Fig. 7). When receiving repeated stimuli with short intervals, the neurons exhibited an incremental increase in response magnitude up to the second stimuli, and then a reduced response in later stimuli (Fig. 2j, l). In contrast, when using long inter-stimulus intervals, the response magnitude was reduced after each stimulus (Fig. 2k, m). These results are consistent with previous observations that habituation is dependent on inter-stimulus durations^20^. Thus, these experiments demonstrate how simple changes of operational parameters allow us to use the same device for a wider repertoire of the device utility.

In contrast to gluing protocols, our system allows for automated imaging by streamlining the handling of the worms; this in turn allows for high-throughput experiments that were not practicable before. To demonstrate the ability to perform rapid screens, we examined the effect of small molecules from an orphan ligand library on mechanosensation. We exposed animals to the compounds in L4 stage, and imaged AVM activity when delivering an anterior stimulus to adult worms (Fig. 3a). Figure 3b shows a typical response of wildtype animals without drug perturbation: calcium traces typically reach a maximum value shortly after the end of stimulus, and then slowly decline back to baseline levels. In order to examine how each drug affects mechanosensation, we quantitatively compared three metrics (max ΔR/Ro, delay time, and half-life), as well as fraction of animals responding, between drug-treated animals and untreated animals (Fig. 3c-h and Supplementary Fig. 8, 9). We imaged adult animals exposed to 13 different drugs and quantified the established parameters for the screen criteria (Supplementary Table 1). While most of the drugs screened lowered the number of animals responding to mechanical stimulus, interestingly, a few drugs slightly increased the response fraction (Fig. 3c). We also analyzed differences in the metrics measure calcium dynamics for the drug treatment conditions, and found that five of the drugs we tested significantly affected mechanosensation response dynamics (Fig. 3d-f). Specifically, D-Alanine (#5) and D-Arginine (#8) significantly attenuated the max ΔR/Ro while increasing the delay in the responses (Fig. 3d, e, h). In contrast, D-Lysine (#4) only attenuated the max response (Fig. 3d, g). Two other drugs had effects only in the decay-time of response (Fig. 3f-h). D-Isoleucine (#3) induced considerably smaller half-life for the calcium transients to return to baseline, much faster than in untreated animals (Fig. 3f, g). Lastly, β-Alanine (#6) significantly increased the half-life, and calcium transients decreased in a slow linear gradient, instead of a typical exponential decay (Fig. 3f, h).

**Figure 3:**
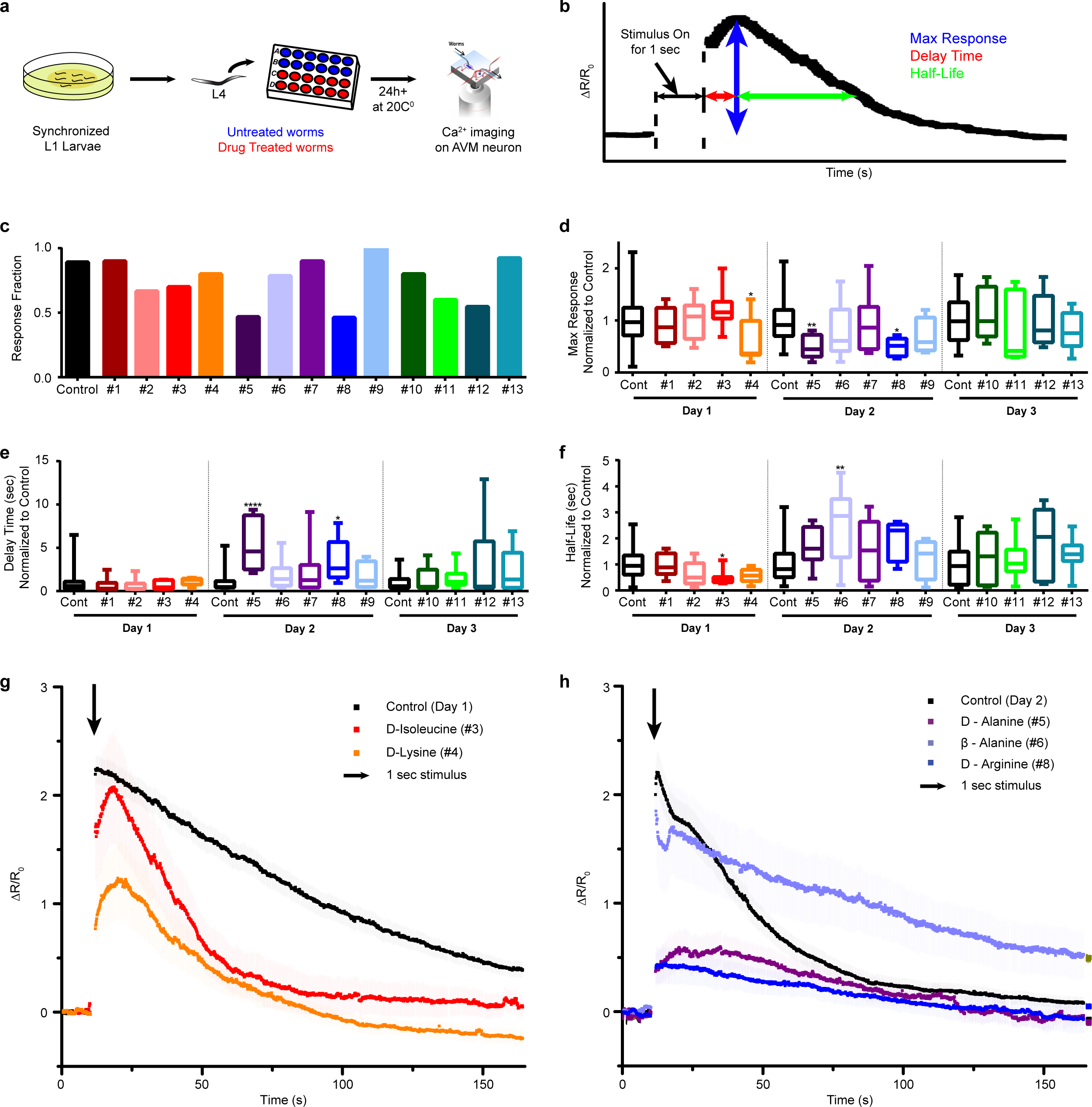
The microfluidic platform enables screens to examine compounds that may affect neuronal responses to mechanical stimuli. **a)** Experimental procedure for the drug screen performed. Synchronized L1 worms are grown in NGM plates to the L4 stage and then deposited in a 48-well plate. Drug treated worms are cultured with 0.5 ml OP50 *E. coli* bacteria (OD 5) and 100μM drugs. Control worms are cultured with 0.5 ml OP50 *E. coli* bacteria (OD 5). Both groups of worms are incubated at 20℃ for at least 24h. Subsequently, AVM responses to 1s stimulus were measured on-chip. **b)** Three metrics measured from individual calcium dynamic traces: maximum response, delay time (time between the end of the stimulus and the arrival of maximum response), and half-life (time it takes the response to decay to half of the maximum). **c)** Fraction of animal responses upon compound treatment. Several compounds produced a lowered fraction of responding animals, while a few slightly increased the response fraction. **d - f)** Box plots show how compounds affect specific parameters of neuronal response upon mechanical stimulation. Quantification of each response was normalized to that of the control group from the same day (day 1 adult to day 3 adult). **d)** D-Lysine (#4), D-Alanine (#5), and D-Arginine (#8) were shown to reduce maximum response, **e)** D-Alanine (#5) and D-Arginine (#8) were shown to increase the time to peak, and **f)** D-Isoluecine (#3) was shown to decrease the decay half-life. In contrast, β-Alanine (#6) was shown to increase the half-life. (Kruskal-Wallis Test, * p<0.05, ** p<0.01, *** p<0.001, **** p<0.0001) **g - h)** Average traces of GCaMP6 in AVM neuron for drug treated worms that cause significant differences from untreated worms in responses to 1s mechanical stimulus. **g)** Day 1 adult worms (Control Day 1: n=53, D-Isoleucine: n=10, D-Lysine: n=10), **h)** Day 2 adult worms (Control Day 2: n=53, D-Alanine: n=15, β-Alanine: n=14, D-Arginine: n=13). Error bar represent SEM.

Another advantage of using microfluidics to deliver mechanosensory stimuli is that it is compatible with other microfluidic components to provoke additional sensory responses, e.g. chemosensation^25^. *C. elegans* is a convenient system for studying multimodal sensory integration *in vivo*; worms have distinct sensory modalities such as mechanosensation and chemosensation, which allow them to find food sources and estimate danger. The difficulty to study sensory integration thus far is that there has not been a convenient method to integrate mechanosensory input with the existing tools, including microfluidic and optical methods^23,25,36–38^. With our mechanical stimulus device, incorporating chemosensory modules is readily attainable by simply adding channels to deliver chemical stimuli (Supplementary Fig. 10); without mechanical stimulation, the response of a chemosensory neuron to a chemical cue is as expected (Supplementary Fig. 11). To demonstrate the utility of the system for multimodal sensory integration, we focused on the response of the PVC command interneurons to both mechanical and chemical stimuli. PVC interneurons are postsynaptic to both the posterior mechanoreceptor neurons PLML/R, as well as the posterior chemosensory neurons PHBL/R, that have been shown to respond to 0.1% SDS stimulus^5,39–41^ (Fig. 4a and Supplementary Fig. 12a). Using our device to deliver multi-modal stimuli, we tested the ability of PVC to respond to stimuli within the same mode and cross-modality. When a single sub-threshold stimulus in either modality is delivered (i.e. 30s SDS stimulation to the tail, or 1s mechanical stimulation to the tail), PVC shows a low probability of response (Fig. 4b, c). Compared to upstream sensory neurons, PVC also responds with a lower magnitude and the response is more variable (Fig. 4b, c and Supplementary Fig. 12b, c). Perhaps not so surprisingly, PVC’s response to subthreshold stimuli in the same modality can be sensitized for subsequent stimulation (Supplementary Fig. 12d). More interestingly, when pre-sensitized by cross-modality sub-threshold stimuli (i.e. chemical before mechanical stimulus, or vice versa), PVC shows similar sensitized responses (Fig. 4d, e, and Supplementary Movie 5, 6). This sensitization is seen both in terms of the magnitude of the individual responses and the fraction of responding animals.

**Figure 4:**
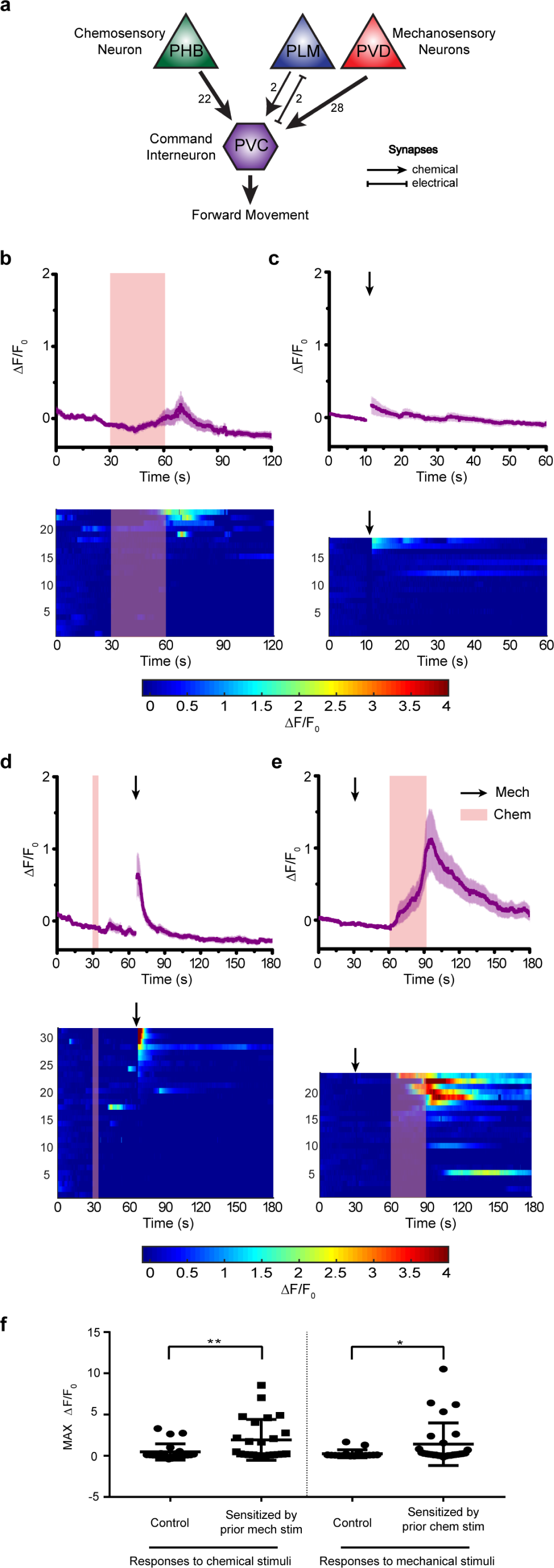
Sensitization of the PVC interneuron responses. **a)** Simplified circuit diagram showing three sensory neurons connecting PVC to forward locomotion behavior (values and arrow thickness indicate number of synapses). **b - c)** Responses of PVC interneuron to a single pulse of stimulation. Averages are plotted on the top graph. Error bars represent SEM. Bottom graphs represent individual traces. **c)** PVC calcium responses to 30s 0.1% SDS stimuli on tail (n=23). **d)** PVC calcium responses to 1s weak mechanical stimuli at 20 psi on posterior region (n=18). In individual traces for outliers, if the value of calcium transient is greater than 4 or less than 0, it would be equal to 4 or 0, respectively (bottom). **d - e)** Sensitized PVC interneuron responses. **e)** Applying 5s 0.1% SDS stimuli enhances the responses of PVC interneuron to 1s weak mechanical stimuli at 20 psi (n=31). Averaged calcium responses (top) and individual traces (bottom) **f)** Applying 1s weak mechanical stimuli at 20 psi enhances the responses of PVC interneuron to 30s 0.1% SDS stimuli (n=24). Averaged calcium responses (top) and individual traces (bottom). Error bars represent SEM (top). In individual traces for outliers, if the value of calcium transient is greater than 4 or less than 0, it would be equal to 4 or 0, respectively (bottom). **f)** Quantified maximum responses of calcium transients to either chemical (left column) or mechanical stimuli (right column). Data points in control groups represent maximum responses to either single chemical or mechanical stimuli. Sensitization of PVC interneuron responses is produced by applying prior weak mechanical stimuli at 20 psi or 5s 0.1% SDS chemical stimuli (Mann-Whitney Test, * p<0.05, ** p<0.01).

## Discussion

For fundamental studies of mechanosensation, quantitative live imaging is necessary, and to perform screens based on mechanosensory phenotypes requires large sample sizes. Our microfluidic platform allows for studying mechanosensation in *C. elegans* quantitatively and conveniently, allowing for the delivery of a variety of types of mechanical stimuli to live animals while recording neuronal activity. Experimental preparation (mainly washing) can be accomplished for a batch of animals, so the limiting step is imaging (tens of seconds to minutes depending on the experiments). Experimental throughput using our streamlined microfluidic system can be as high as ~100 trials per hour; it is also straightforward to automate and run these systems in parallel to further improve throughput. In contrast, the conventional approach (gluing worms and stimulating with a micro stylus and micromanipulator) generally yields ~10successful trials per day. The integration of hardware and software also allows for automated operations of imaging, stimulation, and quantitative analysis, further reducing potential human error and bias. This important improvement in throughput and standardization over conventional methods allowed us to conduct a novel drug screen based on neuronal dynamics due to mechanical stimuli. By using our system, we identified several candidates that strongly affect dynamics in mechanosensory neurons in a variety of ways. One can envision genetic screens performed in a similar manner to identify mechanosensory mutants. Many worm mechanosensory modalities, such as harsh touch and nose touch, involve multiple partially-redundant cell types, making behavioral assays ineffective for finding genes affecting these processes. With simple integration of sorting mechanisms on chip^36,42,43^, it will be possible to conduct high-throughput forward screens for mutants affecting the responses of individual neurons, using a GECI-based assay. The genes identified in such screens should provide insight into the underlying mechanisms of mechanosensation, as well as find potential therapies for sensory-loss conditions such as deafness.

Additionally, microfluidic incorporation of fluidic control can easily allow interrogation of other sensory modalities (e.g. olfaction) in combination with mechanosensation. We have shown that our platform is compatible with previous techniques for delivering chemical stimuli, enabling for the interrogation of integration of multimodal stimuli in the interneurons. This feature can greatly expand the repertoire of assay conditions to allow studies of sensory integration, arousal, habituation, and sensitization. For example, it has been previously shown that neural responses to sensory stimuli become more deterministic as information flows from sensory neurons to interneurons; behavioral responses, however, correlate more strongly with interneurons such as PVC^33,44^. We show here that PVC’s response can be modulated, by prior sensory inputs, and that this modulation is cross-modal. This may point to an interesting ecologically relevant strategy for animal behavior, such that the reliability of the escape response depends both on the stimulus and on the current state of the circuit, as influenced by experience.

Because our platform employs a simple microfluidic device, it is easily adaptable to study biological systems of various sizes. Scaling the devices to be smaller can allow studies of mechanosensory neurons in worm larvae during development; scaling the devices larger can allow studies of the mechanosensation circuit during aging in *C. elegans*, as well as neurons and circuits in other model organisms such as zebrafish or fly larvae. Lastly, because the microfluidic chip allows unhindered optical access, integrations of optogenetic methods^37,38,45–49^ can also be straightforwardly carried out in this platform, thereby greatly expanding the repertoire of biological problems to be studied.

## Acknowledgments

The authors would like to thank K. Shen (HHMI/Stanford) and D. Kim (HHMI Janelia Research Campus) for sharing reagents; GCaMP6 transgene constructs were provided by the Genetically-Encoded Neuronal Indicator and Effector Project, Janelia Research Campus, Howard Hughes Medical Institute. This work was supported in part by the U. S. NIH (R01GM088333 and R01NS096581 to HL), the Wellcome Trust (WT103784MA to WRS), and the MRC (MC-A022-5PB91 to WRS). Some nematode strains used in this work were provided by the Caenorhabditis Genetic Center, which is funded by the NIH, National Center for Research Resources and the International *C. elegans* Knockout Consortium.

### Methods

#### Strains

*C. elegans* were maintained under standard conditions and fed OP50 bacteria^50^. The following strains were used in this study:

AQ 3236 *ljIs142[mec-4::GCaMP6m::SL2TagRFP, unc-119] II; unc-119(ed3) III*

TV17924 *wyls5007[ser2prom3::GCaMP6, egl-17::mCherry] X*

CX10979 *kyEx2865[sra-6::GCaMP3, Pofm-1::GFP]*

GT243 *aEx2[pglr-1::GCaMP6(s), punc-122::GFP]*

RW1596 *stEx30[myo-3p::GFP + rol-6(su1006)]*

To construct AQ3236, we used a single-copy insertion vector containing a GCaMP6M transgene codon-optimized for C. *elegans*, under the control of the *mec-4* promoter (a gift from Doug Kim at HMMI Janelia Research Campus). Single-copy chromosomal integrations were obtained using the MosTic procedure^51^. Unless otherwise specified, all worms imaged in this study are adults.

#### Chip Design and Fabrication

The device consists of worm inlet/outlet, imaging channel (50~60 µm deep), and four sets of actuated PDMS membranes. Animals loosely fit in the channel, and are trapped (but not held) in the imaging area by two sets of actuated members. The width of actuated PDMS membrane is 150 µm, the distance between first and second sets of membrane is 200 µm and second and third sets of membrane is 250 µm.

Since worms were not immobilized using drugs, animals’ head or tail can move in the imaging channel of the microfluidic chip. This movement sometimes blurs images. To reduce the movement of head or tail part of worms, a three-step vertical tapering of the imaging channel was used to restrict the out-of-plane movement. The thickness of first and second layers was 15 µm and third layer was 20 µm for the 50 µm deep imaging channel; these layers were created by SU-8 2015 negative photoresist (MicroChem) using standard photolithographic techniques^52^.

To create the actuated PDMS structure to touch and trap worms, multi-layer soft lithography process^53^ was used. For the bottom flow layer of features, 23:1 PDMS was deposited via spin coating to create a thin layer. For the top control layer, 10:1 PDMS was directly poured onto a blank master, which does not have any features, to create a thick and mechanically rigid handle layer. Both layers were then placed into a 90°C oven for 25-30 minutes until the control layer PDMS was rigid but sticky. After they were manually aligned, additional 10:1 PDMS was poured and cured for several hours to create a rigid handling layer for the device.

#### Calcium Imaging

All imaging experiments were performed on a Leica DMIRB inverted microscope using a 40x air objective (N. A. 0.75). Video sequences were captured using a Hamamatsu EM-CCD camera with 100 ms exposure time. Simultaneous two-color imaging was performed using a DV2 beamsplitter (Photometrics) containing a GFP/RFP filter set. Excitation light for fluorescent imaging was delivered through a projector system previously developed^38^. In experiments for the measurement of mechanosensory neuronal responses, stimuli were delivered 10 s after recording baseline activity of neurons. In experiment for the measurement of interneuronal recording, stimuli were delivered 30 s after recording baseline activity of neurons. Videos were recorded for 60-180 s following stimulus delivery.

#### Data Analysis

Fluorescence intensities for each frame were extracted using customized neuron-tracking Matlab scripts (Supplementary Fig. 7). In strains where both GCaMP6 and RFP are expressed, the ratio between intensity values were computed 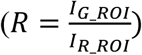 in order to minimize movement artifacts. When only GCaMP was available, fluorescence values were computed by subtracting background intensity 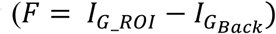. GCaMP and RFP intensities were measured as the mean pixel intensity of the 100 brightest pixels of a circular region of interest (ROI) of 10 pixel radius. Background intensities were subtracted to adjust for variations in lighting conditions, and were measured as the mean pixel intensity of an ROI in a background region (Supplementary Fig. 7). Calcium traces were computed as the change in R or F from the baseline value 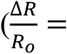 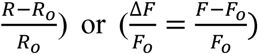. Baseline values were computed as the mean R or F prior to stimulus delivery.

#### Drug screening

Worms were roughly synchronized by picking 20-25 L4 worms and allowing them to lay eggs overnight before removing them from the plate. After two days at 20°C, tightly age-synchronized populations of worms were obtained by washing adults and L1s off of these plates and then washing newly hatched L1s from these plates after an hour interval. The 84 compounds of the Screen-Well Orphan library (ENZO) were used for the drug screening. 20-30 tightly-synchronized L4 worms were placed on a 48-well plate (Greiner Bio-One) with 0.5 ml OP50 bacteria (OD 5) for non-treated worms and both 0.495 ml OP50 bacteria and 0.005 ml (100 µM) drugs for drug-treated worms. After 24 hours, worms were imaged. Among 84 compounds in the library, we tested the effect of 13 compounds on AVM neuronal responses at three different ages (from day 1 adult to day 3 adult). These compounds were chosen randomly from the orphan ligand library.

---

**Supplementary Figure 1:**
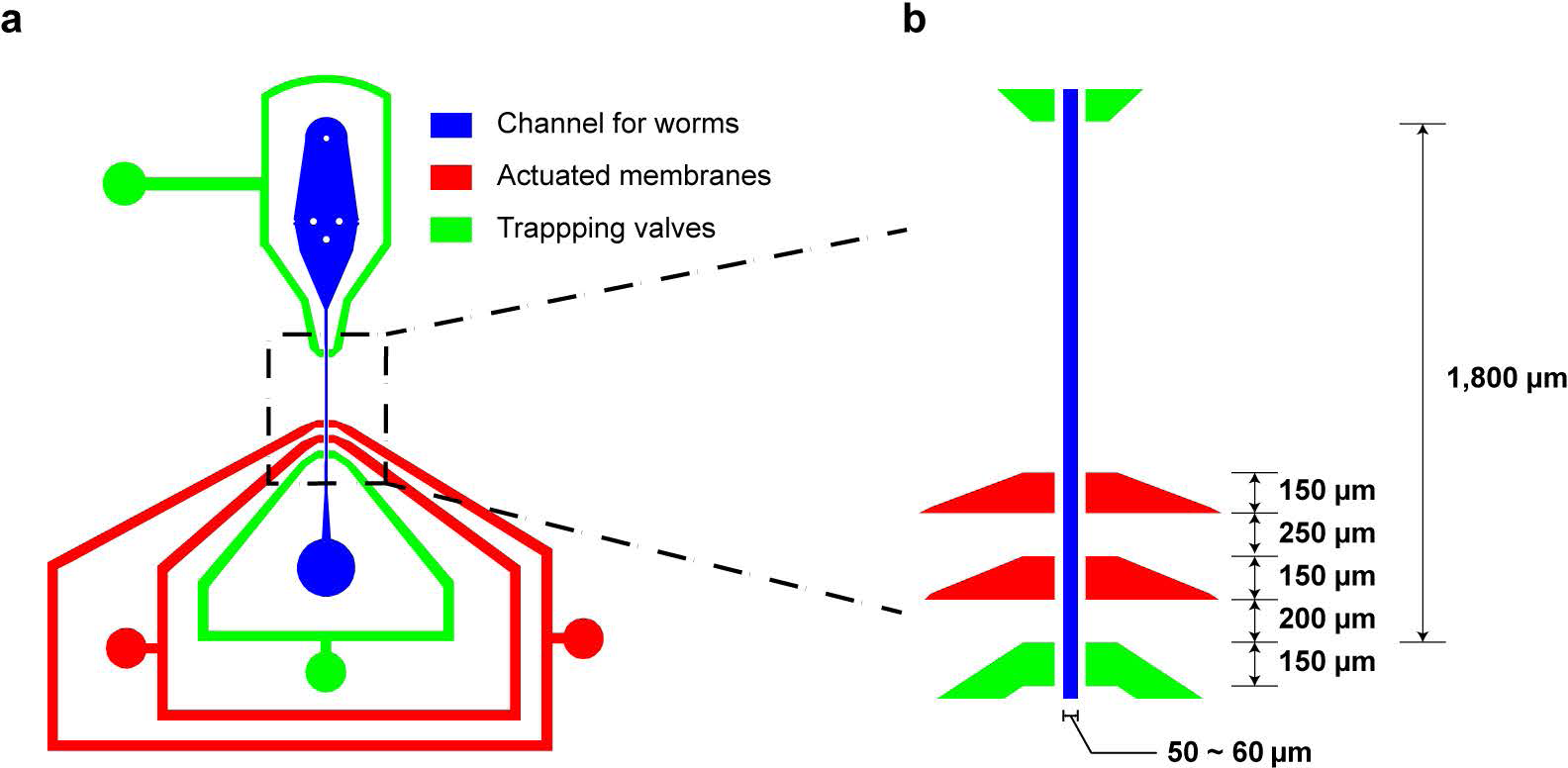
Overview of microfluidic device design and dimensions. The device is composed of the channel for worms (50-60 μm deep and wide, which allow the animals to fit loosely inside), two sets of actuated membrane, and two sets of trapping valve. The width of the both of actuated PDMS membrane and trapping valve is 150 μm, the distance between first and second sets of membrane is 200 μm and second and third sets of membrane is 250 μm.

**Supplementary Figure 2:**
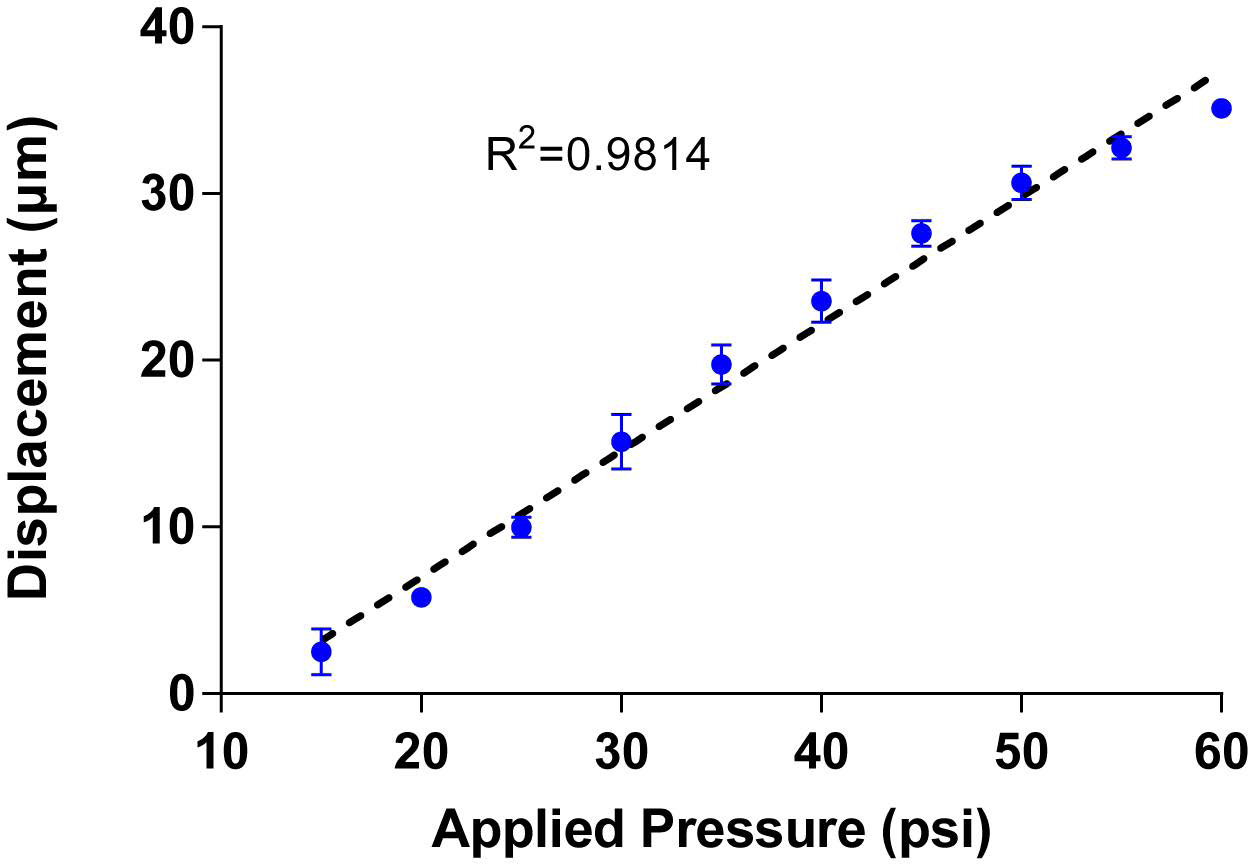
Displacement of the actuated membrane by applying pressure (n=4worms). It is important to note that the measurements were taken by images from transgenic worms expressing GFP along body-wall muscle (*stEx30[myo-3p::GFP + rol-6(su1006)]*). R-square value is 0.9814.

**Supplementary Figure 3:**
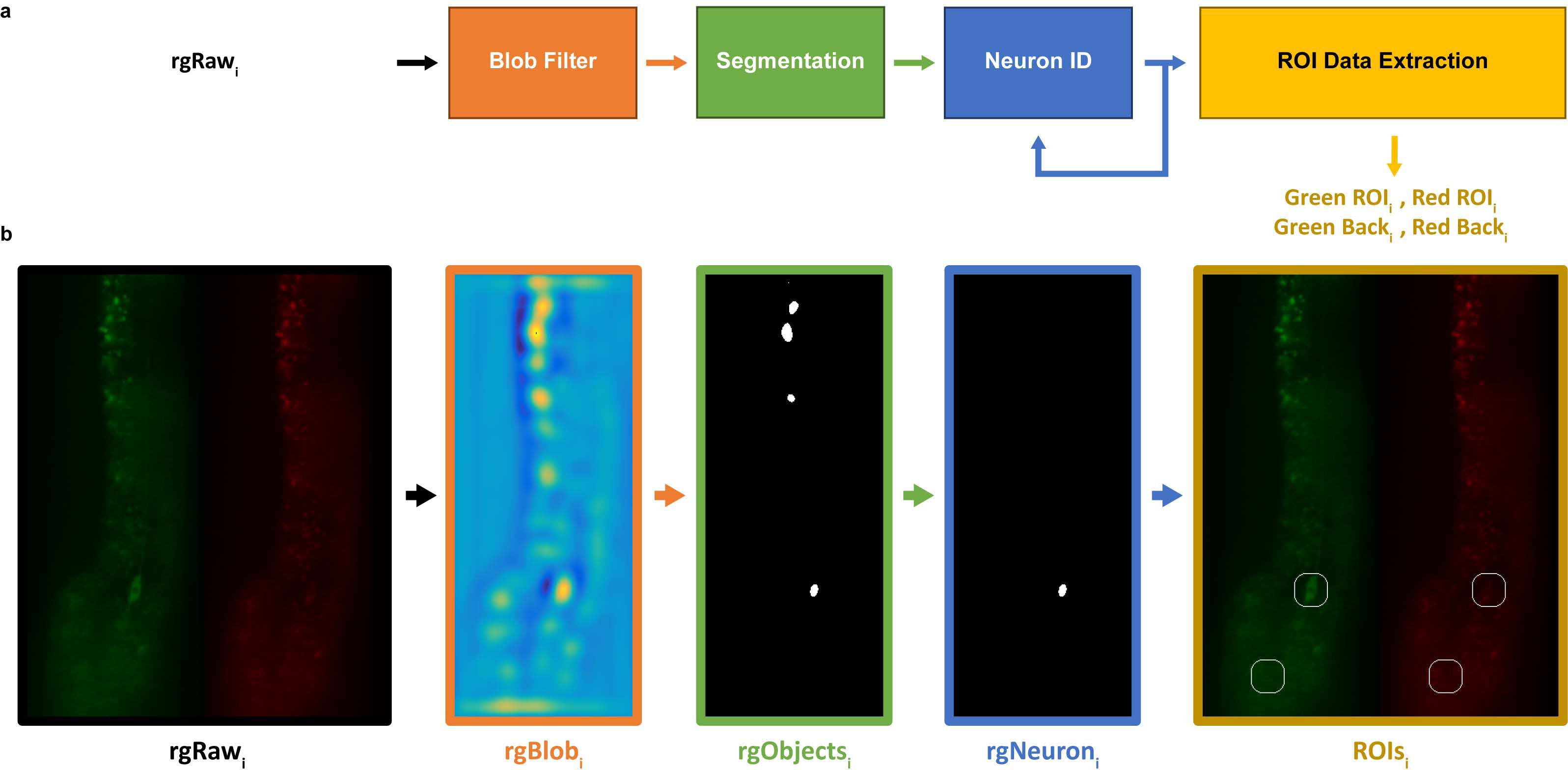
Neuron Tracking Algorithm. In order to extract fluorescence intensities throughout recordings, a neuron tracking algorithm was developed. This was necessary because worms are not fully immobilized in the device, and mechanical stimuli often caused the neuron of interest to move within the field-of-view. **a)** Overall schematic of the neuron tracking algorithm. For each frame *i*, raw images are processed through a blob filter (Laplace of Gaussian filter) to improve contrast and facilitate segmentation. Blob filtered images are segmented by applying an empirically determined threshold. The neuron of interest is identified by the user in the first frame, and by distance to the neuron in the previous frame. Lastly, once the neuron is detected for each frame, intensity values are extracted (Green ROI, Red ROI, Green Background, and Red Background). **b)** Example of algorithm procedure.

**Supplementary Figure 4:**
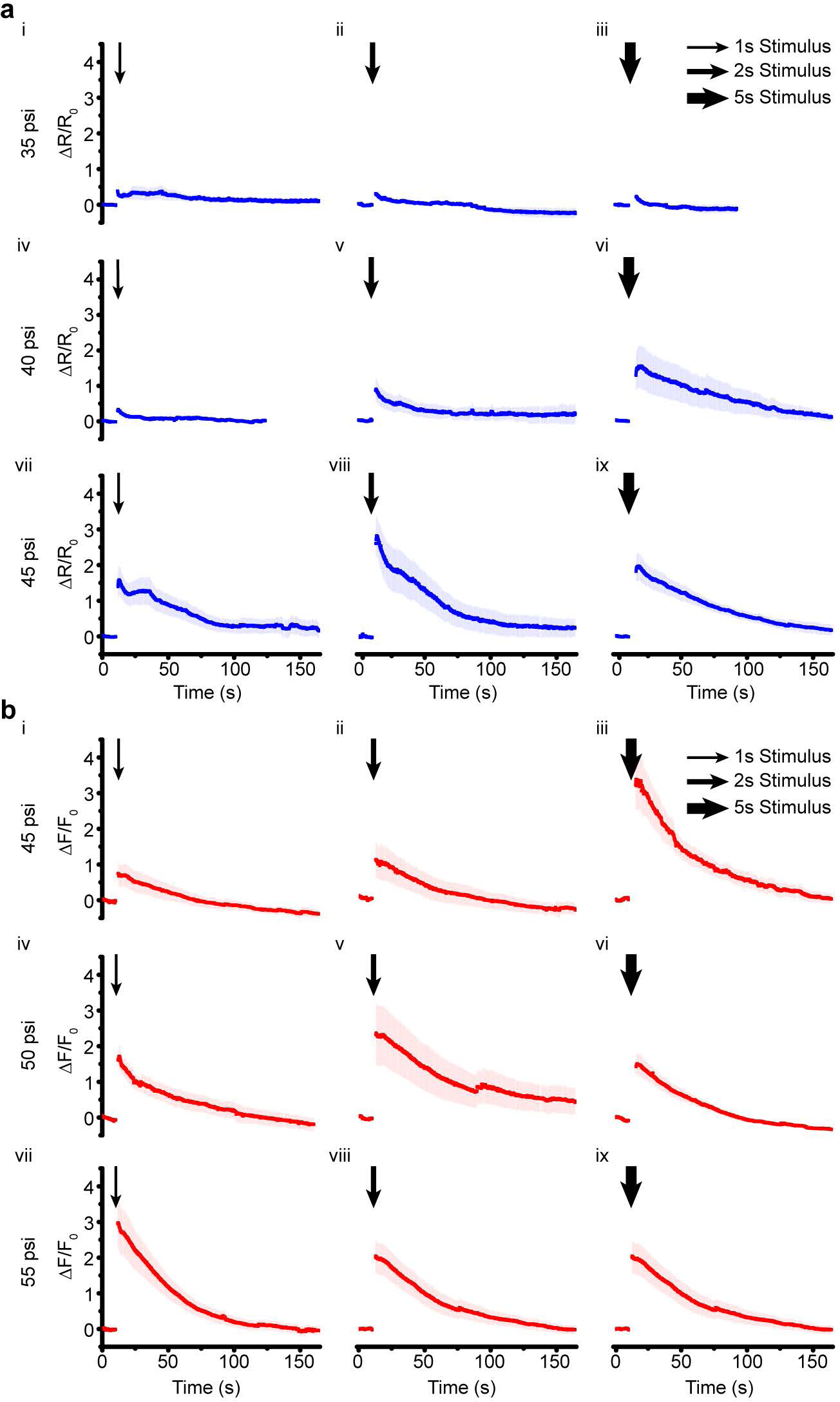
**a)** Average traces of GCaMP6 signal in AVM neuron in response to diverse pressures and stimulus durations (**i-iii:** 35 psi, **iv-vi:** 40 psi, **vii-ix:** 45 psi / **i, iv, vii:** 1s stimulus, **ii, v, viii:** 2s stimulus, **iii, vi, ix:** 5s stimulus, sample size **i:** n=25, **ii:** n=10, **iii:** n=8, **iv:** n=8, **v:** n=10, **vi:** n=10, **vii:** n=27, **viii:** n=6, **ix:** n=10). Error bars represent SEM. **b)** Average traces of GCaMP6 signal in PVD neuron in response to diverse pressures and stimulus durations (**i-iii:** 45 psi, **iv-vi:** 50 psi, **vii-ix:** 55 psi / **i, iv, vii:** 1s stimulus, **ii, v, viii:** 2s stimulus, **iii, vi, ix:** 5s stimulus, sample size **i:** n=9, **ii:** n=4, **iii:** n=6, **iv:** n=6, **v:** n=9, **vi:** n=10, **vii:** n=9, **viii:** n=10, **ix:** n=10). Error bar represent SEM.

**Supplementary Figure 5:**
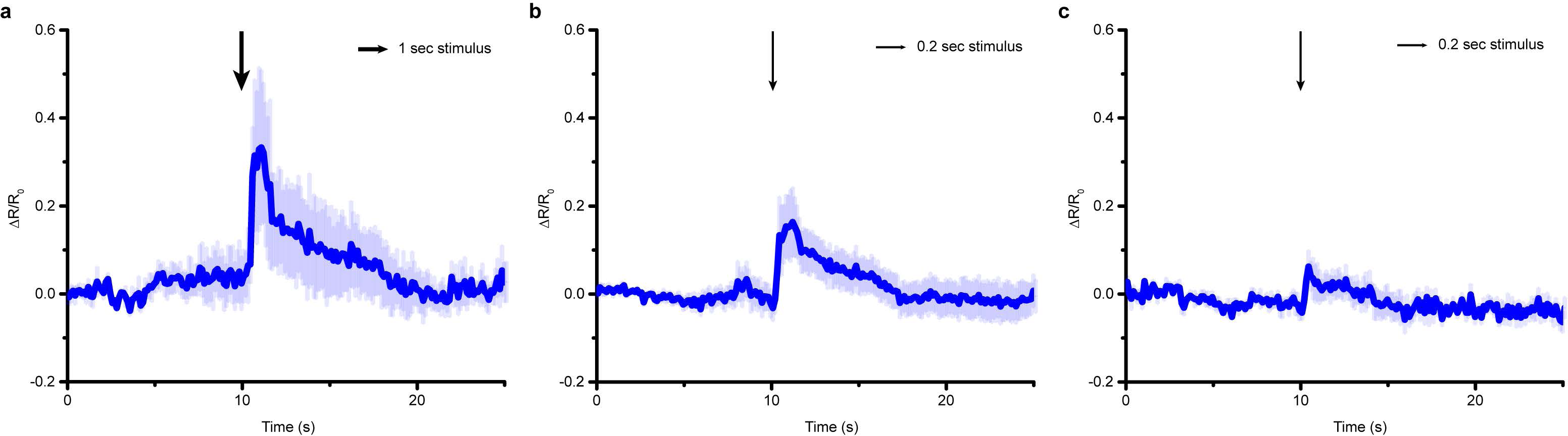
AVM cell body responses to various stimuli with low pressures and durations **a)** 30 psi and 1 s (n=4), **b)** 30 psi and 0.2 s (n=10), **c)** 15 psi and 0.2 s (n=5). AVM response is reduced when using lower pressures (comparing a to Fig. 2a, and a to c). Response is also attenuated when using shorter durations (comparing a to b). Error bars represent SEM.

**Supplementary Figure 6:**
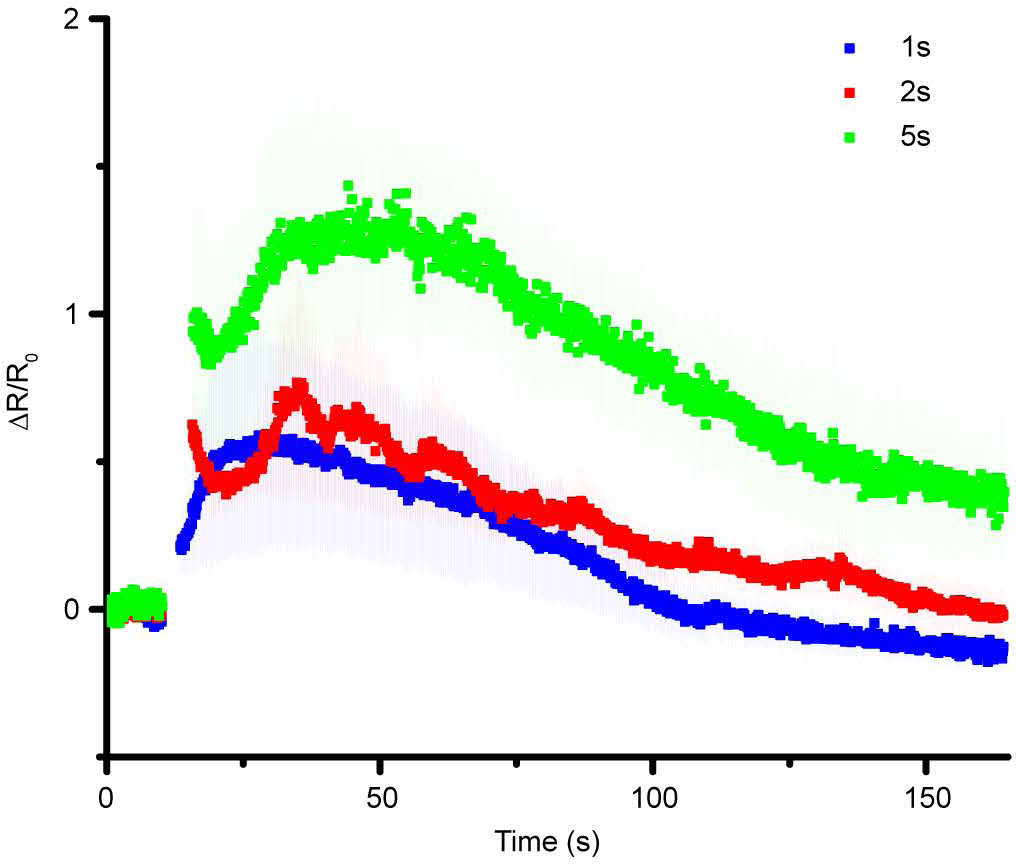
PLM cell body responses to various stimulus durations (1s: n=9, 2s: n=4, 5s: n=4). Similar to those of AVM, maximum responses in PLM were proportional to the stimulus duration.

**Supplementary Figure 7:**
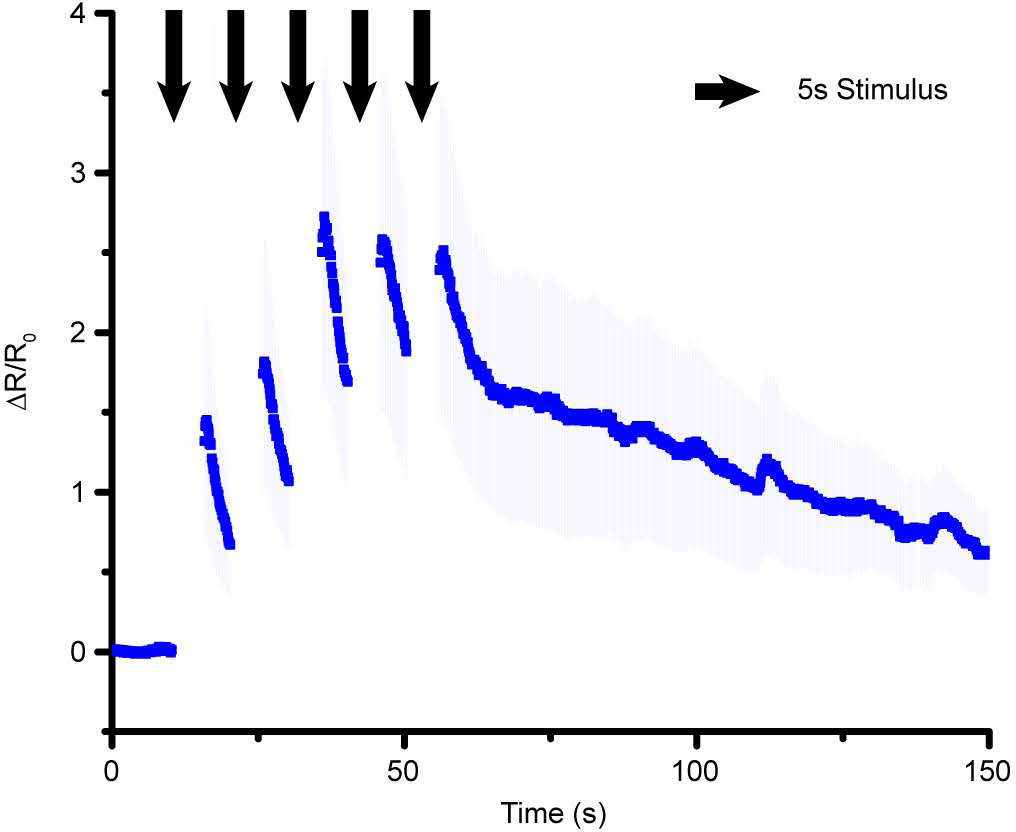
AVM cell body response to delivery of repeated stimuli with long durations (5s, n=5). Similar to Figure 2l, traces showed incremental increases in the first few stimuli, and showed a decreased response in later stimuli.

**Supplementary Figure 8:**
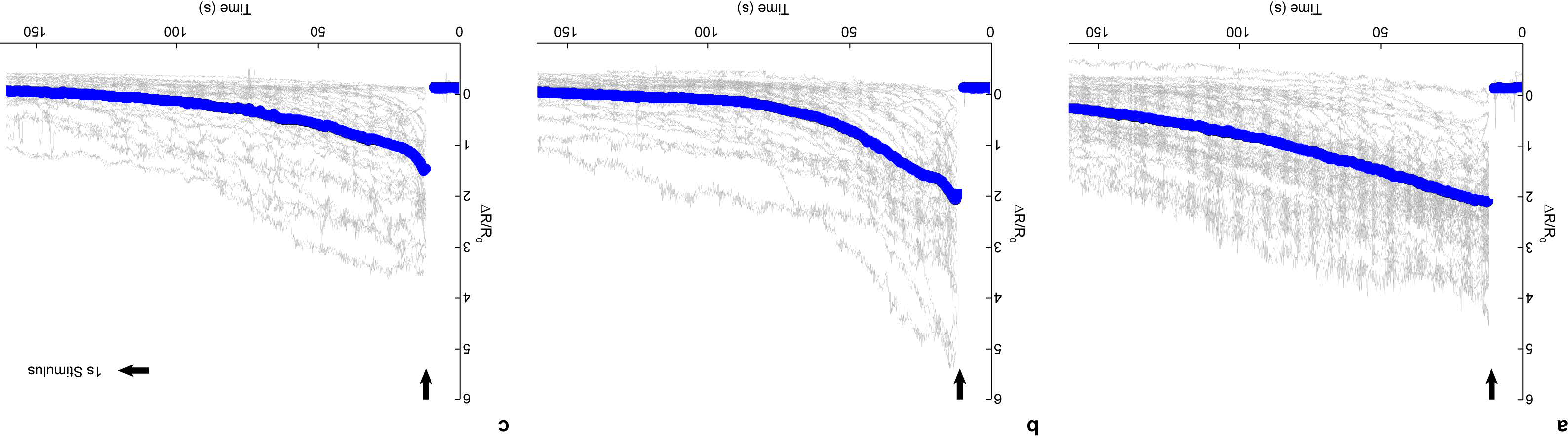
Individual (gray) and average traces (blue) for AVM response in untreated animals for different control groups for drug screen. **a)** Day 1 (n=53), **b)** Day 2 (n=53), **c)** Day 3 (n=35) adult worms.

**Supplementary Figure 9:**
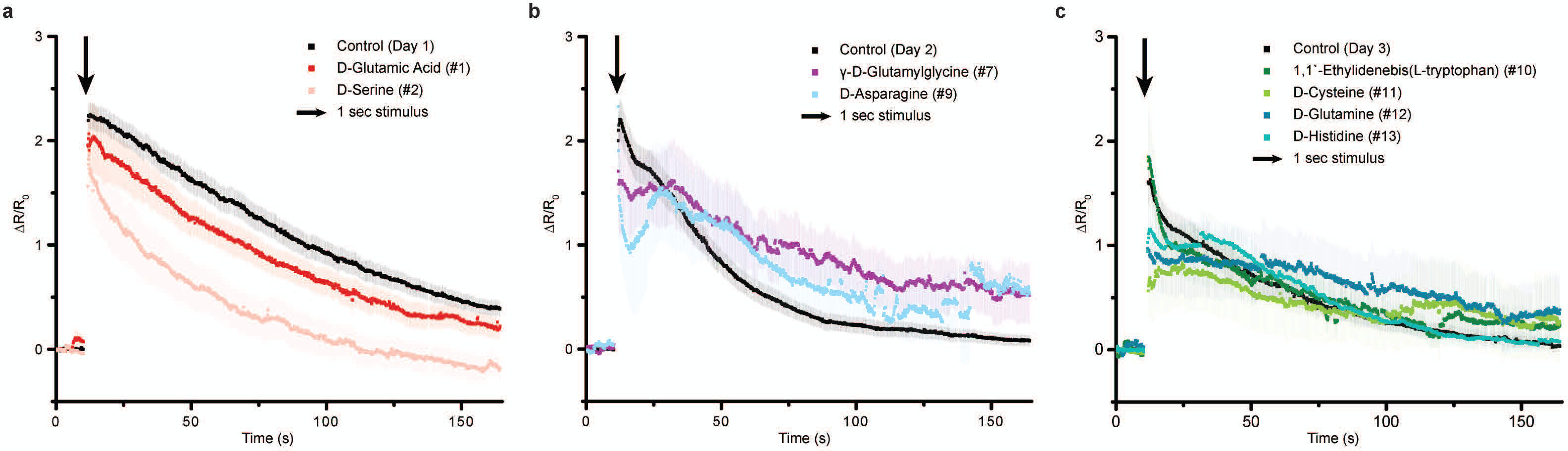
Average traces for AVM response in drug-treated animals that do not show a significant difference from the control groups. **a)** Day 1 adult worms (Control Day 1: n=53, D-Glutamic acid: n=10, D-Serine: n=12), **b)** Day 2 adult worms (Control Day 2: n=53, γ-D-Glutamylglycine: n=10, D-Asparagine: n=4), **c)** Day 3 adult worms (Control Day 3: n=35, 1,1'-Ethylidene-bis(L-tryptophan): n=10, D-Cysteine: n=10, D-Glutamine: n=11, D-Histidine: n=13).

**Supplementary Figure 10:**
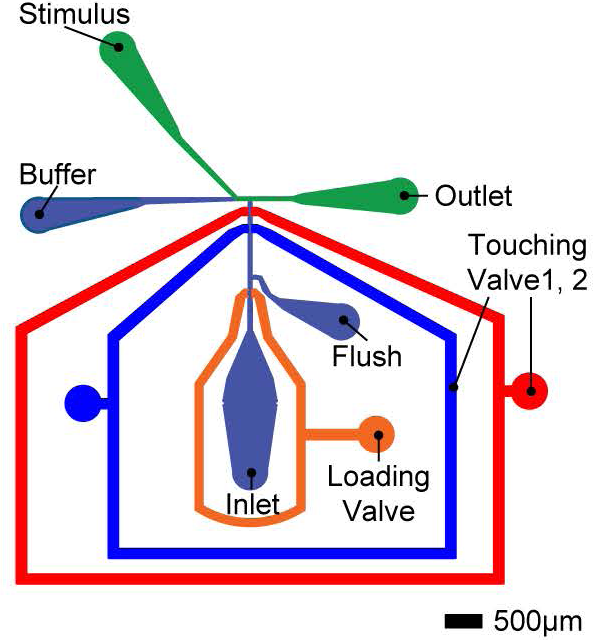
Overview of microfluidic device for the delivery of multimodal stimuli. The device is composed of a channel for worms (Inlet and Flush channel), two sets of actuated membrane for mechanical stimuli (Touching valve 1,2), one set of trapping valve (Loading valve), two inlets for chemical stimuli (Buffer and Stimulus), and outlet.

**Supplementary Figure 11:**
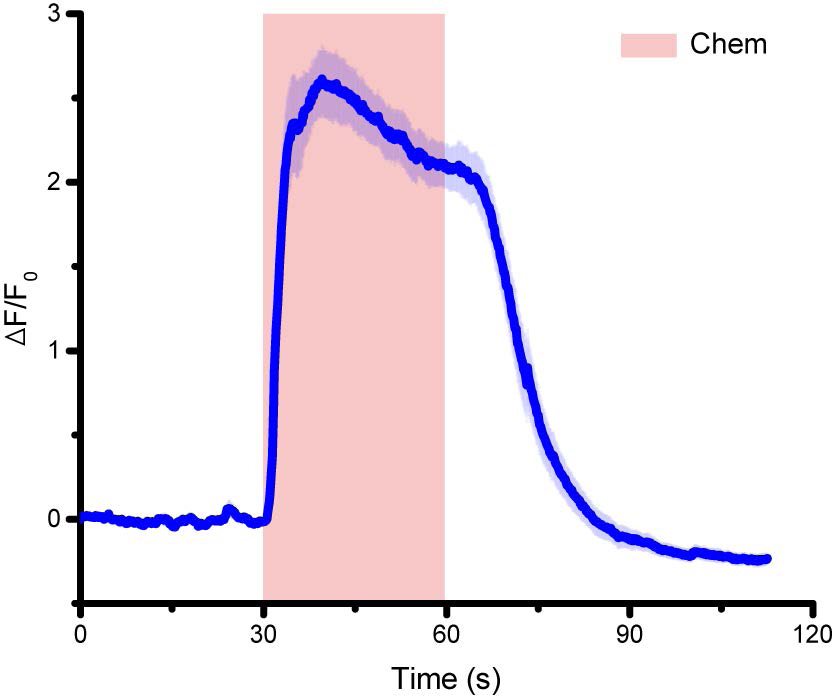
Average traces of GCaMP3 signal in ASH neuron in response to 30s 0.1% SDS stimuli (n=13). Stimuli were delivered 30 s after recording baseline activity of neurons. Error bars represent SEM.

**Supplementary Figure 12:**
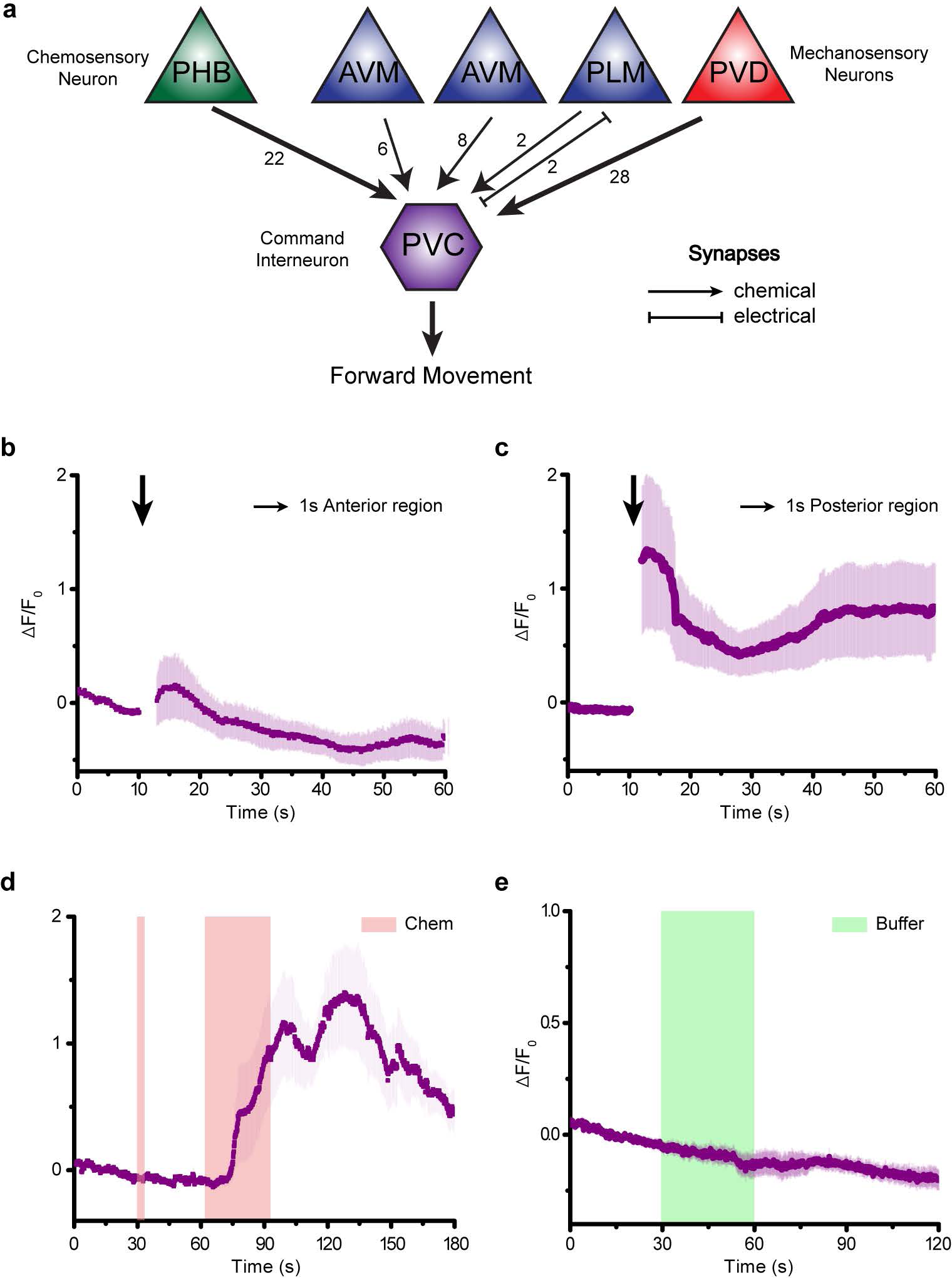
**a)** Neural wiring diagram showing five sensory neurons in a circuit linking PVC to forward behavior, and the number of direct synapses between each pair of neurons (arrow thickness indicates number of synapses). **b-c)** The activity of PVC responses to localized strong mechanical stimuli: **b)** 1s anterior stimuli (n=5) and **c)** 1s posterior stimuli (n=18) at 45psi. **d)** Applying prior 5s 0.1% SDS stimuli enhances the responses of PVC interneuron to next 30s 0.1% SDS stimuli (n=13). **e)** The activity of PVC responses to buffer to buffer changes (n=10). Error bars represents SEM.

**Supplementary Table 1:** The 13 compounds of the orphan library were used for the drug screen (Fig. 3). Sample size is the total number of tested worms and if the value of maximum responses is larger than 0.5, it is counted as a responding worm.

| Number | Name | Rationale | Sample size | Number of responding worms |
| --- | --- | --- | --- | --- |
| 1 | D-Glutamic acid | Putative endogenous ligand | 10 | 9 |
| 2 | D-Serine | Putative endogenous ligand | 12 | 8 |
| 3 | D-Isoleucine | D-Amino acid | 10 | 7 |
| 4 | D-Lysine | D-Amino acid | 10 | 8 |
| 5 | D-Alanine | D-Amino acid | 15 | 7 |
| 6 | β-Alanine | Endogenous | 14 | 11 |
| 7 | γ-D-Glutamylglycine | D-Amino acid | 10 | 9 |
| 8 | D-Arginine | D-Amino acid | 13 | 6 |
| 9 | D-Asparagine | D-Amino acid | 4 | 4 |
| 10 | 1,1'-Ethylidene-bis(L-tryptophan) | Bioactive tryptophan derivative | 10 | 8 |
| 11 | D-Cysteine | D-Amino acid | 10 | 6 |
| 12 | D-Glutamine | D-Amino acid | 11 | 6 |
| 13 | D-Histidine | D-Amino acid | 13 | 12 |

**Supplementary Movie 1**: Calcium dynamics of AVM neuron to 1s anterior stimulation at 45psi. Stimulus was delivered 10s after recording baseline of neuronal activity. The transgenic animal shown here expresses GCaMP6 and RFP in AVM neuron (*ljIs142[mec-4::GCaMP6m::SL2TagRFP, unc-119] II; unc-119(ed3) III*). Left panel shows green fluorescence from GCaMP6m (left) and red fluorescence from RFP (right) in false colors. White boxes indicate location of AVM neuron and shows how algorithm tracks the neuron. Right graph shows the quantitative calcium trance and red circle indicates the current time point of video. Stimulus occurs at 10s (red dash line). 1x playback.

**Supplementary Movie 2**: Calcium dynamics of PLM neuron to 1s posterior stimulation at 45psi. Stimulus was delivered 10s after recording baseline of neuronal activity. The transgenic animal shown here expresses GCaMP6 and RFP in PLM neuron (*ljIs142[mec-4::GCaMP6m::SL2TagRFP, unc-119] II; unc-119(ed3) III*). Left panel shows green fluorescence from GCaMP6m (left) and red fluorescence from RFP (right) in false colors. White boxes indicate location of PLM neuron and shows how algorithm tracks the neuron. Right graph shows the quantitative calcium trance and red circle indicates the current time point of video. Stimulus occurs at 10s (red dash line). 1x playback.

**Supplementary Movie 3**: Calcium dynamics of PVM neuron to 1s posterior stimulation at 45psi. Stimulus was delivered 10s after recording baseline of neuronal activity. The transgenic animal shown here expresses GCaMP6 and RFP in PVM neuron (*ljIs142[mec-4::GCaMP6m::SL2TagRFP, unc-119] II; unc-119(ed3) III*). Left panel shows green fluorescence from GCaMP6m (left) and red fluorescence from RFP (right) in false colors. White boxes indicate location of PVM neuron and shows how algorithm tracks the neuron. Right graph shows the quantitative calcium trance and red circle indicates the current time point of video. Stimulus occurs at 10s (red dash line). 1x playback.

**Supplementary Movie 4**: Calcium dynamics of PVD neuron to 1s posterior stimulation at 45psi. Stimulus was delivered 10s after recording baseline of neuronal activity. The transgenic animal shown here expresses GCaMP6 in PVD neuron (*wyls5007[ser2prom3::GCaMP6, egl-17::mCherry] X*). Left panel shows green fluorescence from GCaMP6 in false color. A white box indicates location of PVD neuron and shows how algorithm tracks the neuron. Right graph shows the quantitative calcium trance and red circle indicates the current time point of video. Stimulus occurs at 10s (red dash line). 5x playback.

**Supplementary Movie 5**: Calcium dynamics of PVC interneuron. Applying 5s 0.1% SDS chemical stimulus at 30s (red dash lines on the right panel) and then 1s weak mechanical stimuli at 65s (blue dash lines on the right panel). The transgenic animal shown here expresses GCaMP6 in PVC interneuron (*aEx2[pglr-1::GCaMP6(s), punc-122::GFP]*). Left panel shows green fluorescence from GCaMP6 in false color. A white box indicates location of PVC interneuron and shows how algorithm tracks the neuron. Right graph shows the quantitative calcium trance and red circle indicates the current time point of video. Stimulus occurs at 10s (red dash line). 5x playback.

**Supplementary Movie 6**: Calcium dynamics of PVC interneuron. Applying 1s weak mechanical stimulus at 30s (blue dash lines on the right panel) and then 30s 0.1% SDS chemical stimulus at 60s (red dash lines on the right panel). The transgenic animal shown here expresses GCaMP6 in PVC interneuron (*aEx2[pglr-1::GCaMP6(s), punc-122::GFP]*). Left panel shows green fluorescence from GCaMP6 in false color. A white box indicates location of PVC interneuron and shows how algorithm tracks the neuron. Right graph shows the quantitative calcium trance and red circle indicates the current time point of video. Stimulus occurs at 10s (red dash line). 5x playback.

## References

1 Corey, D. P. New TRP channels in hearing and mechanosensation. Neuron 39, 585–588 (2003).

2 Kung, C. A possible unifying principle for mechanosensation. Nature 436, 647–654 (2005).

3 Nicolson, T. The genetics of hearing and balance in zebrafish. Annu. Rev. Genet. 39, 9–22 (2005).

4 Lumpkin, E. A. & Caterina, M. J. Mechanisms of sensory transduction in the skin. Nature 445, 858–865 (2007).

5 Chalfie, M. et al. The neural circuit for touch sensitivity in Caenorhabditis elegans. The Journal of neuroscience 5, 956–964 (1985).

6 Chalfie, M. & Au, M. Genetic control of differentiation of the Caenorhabditis elegans touch receptor neurons. Science 243, 1027–1033 (1989).

7 Hong, K. & Driscoll, M. A transmembrane domain of the putative channel subunit MEC-4 influences mechanotransduction and neurodegeneration in C. elegans. (1994).

8 Huang, M. & Chalfie, M. Gene interactions affecting mechanosensory transduction in Caenorhabditis elegans. Nature 367, 467–470 (1994).

9 Corey, D. P. & Garcia-Anoveros, J. Mechanosensation and the DEG/ENaC ion channels. Science 273, 323 (1996).

10 Tavernarakis, N., Shreffler, W., Wang, S. & Driscoll, M. unc-8, a DEG/ENaC family member, encodes a subunit of a candidate mechanically gated channel that modulates C. elegans locomotion. Neuron 18, 107–119 (1997).

11 Tobin, D. M. et al. Combinatorial expression of TRPV channel proteins defines their sensory functions and subcellular localization in C. elegans neurons. Neuron 35, 307–318 (2002).

12 Goodman, M. B. et al. MEC-2 regulates C. elegans DEG/ENaC channels needed for mechanosensation. Nature 415, 1039–1042 (2002).

13 O'Hagan, R., Chalfie, M. & Goodman, M. B. The MEC-4 DEG/ENaC channel of Caenorhabditis elegans touch receptor neurons transduces mechanical signals. Nature neuroscience 8, 43–50 (2005).

14 Kindt, K. S. et al. Caenorhabditis elegans TRPA-1 functions in mechanosensation. Nature neuroscience 10, 568–577 (2007).

15 Chatzigeorgiou, M. et al. Specific roles for DEG/ENaC and TRP channels in touch and thermosensation in C. elegans nociceptors. Nature neuroscience 13, 861–868 (2010).

16 Delmas, P. & Coste, B. Mechano-gated ion channels in sensory systems. Cell 155, 278–284 (2013).

17 Sulston, J., Dew, M. & Brenner, S. Dopaminergic neurons in the nematode Caenorhabditis elegans. Journal of Comparative Neurology 163, 215–226 (1975).

18 Chalfie, M. & Sulston, J. Developmental genetics of the mechanosensory neurons of Caenorhabditis elegans. Developmental biology 82, 358–370 (1981).

19 Way, J. C. & Chalfie, M. The mec-3 gene of Caenorhabditis elegans requires its own product for maintained expression and is expressed in three neuronal cell types. Genes & development 3, 1823–1833 (1989).

20 Suzuki, H. et al. In vivo imaging of C. elegans mechanosensory neurons demonstrates a specific role for the MEC-4 channel in the process of gentle touch sensation. Neuron 39, 1005–1017 (2003).

21 Eastwood, A. L. et al. Tissue mechanics govern the rapidly adapting and symmetrical response to touch. Proceedings of the National Academy of Sciences 112, E6955–E6963 (2015).

22 Whitesides, G. M. The origins and the future of microfluidics. Nature 442, 368–373 (2006).

23 San-Miguel, A. & Lu, H. Microfluidics as a tool for C. elegans research. (2005).

24 Ryu, W. S. & Samuel, A. D. Thermotaxis in Caenorhabditis elegans analyzed by measuring responses to defined thermal stimuli. The Journal of neuroscience 22, 5727–5733 (2002).

25 Chronis, N., Zimmer, M. & Bargmann, C. I. Microfluidics for in vivo imaging of neuronal and behavioral activity in Caenorhabditis elegans. Nature methods 4, 727–731 (2007).

26 Chalasani, S. H. et al. Dissecting a circuit for olfactory behaviour in Caenorhabditis elegans. Nature 450, 63–70 (2007).

27 Kuhara, A. et al. Temperature sensing by an olfactory neuron in a circuit controlling behavior of C. elegans. Science 320, 803–807 (2008).

28 Guo, Z. V., Hart, A. C. & Ramanathan, S. Optical interrogation of neural circuits in Caenorhabditis elegans. Nature methods 6, 891–896 (2009).

29 Macosko, E. Z. et al. A hub-and-spoke circuit drives pheromone attraction and social behaviour in C. elegans. Nature 458, 1171–1175 (2009).

30 Ha, H.-i. et al. Functional organization of a neural network for aversive olfactory learning in Caenorhabditis elegans. Neuron 68, 1173–1186 (2010).

31 Albrecht, D. R. & Bargmann, C. I. High-content behavioral analysis of Caenorhabditis elegans in precise spatiotemporal chemical environments. Nature methods 8, 599–605 (2011).

32 Kato, S., Xu, Y., Cho, C. E., Abbott, L. & Bargmann, C. I. Temporal responses of C. elegans chemosensory neurons are preserved in behavioral dynamics. Neuron 81, 616–628 (2014).

33 Luo, L. et al. Dynamic encoding of perception, memory, and movement in a C. elegans chemotaxis circuit. Neuron 82, 1115–1128 (2014).

34 Chen, T.-W. et al. Ultrasensitive fluorescent proteins for imaging neuronal activity. Nature 499, 295–300 (2013).

35 Li, W., Kang, L., Piggott, B. J., Feng, Z. & Xu, X. S. The neural circuits and sensory channels mediating harsh touch sensation in C. elegans. Nature communications 2, 315 (2011).

36 Crane, M. M., Chung, K., Stirman, J. & Lu, H. Microfluidics-enabled phenotyping, imaging, and screening of multicellular organisms. Lab on a Chip 10, 1509–1517 (2010).

37 Leifer, A. M., Fang-Yen, C., Gershow, M., Alkema, M. J. & Samuel, A. D. Optogenetic manipulation of neural activity in freely moving Caenorhabditis elegans. Nature methods 8, 147–152 (2011).

38 Stirman, J. N. et al. Real-time multimodal optical control of neurons and muscles in freely behaving Caenorhabditis elegans. Nature methods 8, 153–158 (2011).

39 Hilliard, M. A., Bargmann, C. I. & Bazzicalupo, P. C. elegans responds to chemical repellents by integrating sensory inputs from the head and the tail. Current Biology 12, 730–734 (2002).

40 White, J. G., Southgate, E., Thomson, J. N. & Brenner, S. The structure of the nervous system of the nematode Caenorhabditis elegans. Philos Trans R Soc Lond B Biol Sci 314, 1–340 (1986).

41 Wicks, S. R. & Rankin, C. H. Integration of mechanosensory stimuli in Caenorhabditis elegans. The Journal of neuroscience 15, 2434–2444 (1995).

42 Chung, K., Crane, M. M. & Lu, H. Automated on-chip rapid microscopy, phenotyping and sorting of C. elegans. Nature methods 5, 637–643 (2008).

43 Crane, M. M., Chung, K. & Lu, H. Computer-enhanced high-throughput genetic screens of C. elegans in a microfluidic system. Lab Chip 9, 38–40 (2009).

44 Gordus, A., Pokala, N., Levy, S., Flavell, S. W. & Bargmann, C. I. Feedback from network states generates variability in a probabilistic olfactory circuit. Cell 161, 215–227 (2015).

45 Boyden, E. S., Zhang, F., Bamberg, E., Nagel, G. & Deisseroth, K. Millisecond-timescale, genetically targeted optical control of neural activity. Nature neuroscience 8, 1263–1268 (2005).

46 Zhang, F. et al. Multimodal fast optical interrogation of neural circuitry. Nature 446, 633–639 (2007).

47 Airan, R. D., Thompson, K. R., Fenno, L. E., Bernstein, H. & Deisseroth, K. Temporally precise in vivo control of intracellular signalling. Nature 458, 1025–1029 (2009).

48 Deisseroth, K. Optogenetics. Nature Methods 8, 26–29, doi:10.1038/nmeth.f.324 (2011).

49 Larsch, J. et al. A Circuit for Gradient Climbing in C. elegans Chemotaxis. Cell Rep 12, 1748–1760 (2015).

50 Brenner, S. The genetics of Caenorhabditis elegans. Genetics 77, 71–94 (1974).

51 Frøkjær-Jensen, C. et al. Single-copy insertion of transgenes in Caenorhabditis elegans. Nature genetics 40, 1375–1383 (2008).

52 del Campo, A. & Greiner, C. SU-8: a photoresist for high-aspect-ratio and 3D submicron lithography. Journal of Micromechanics and Microengineering 17, R81 (2007).

53 Unger, M. A., Chou, H.-P., Thorsen, T., Scherer, A. & Quake, S. R. Monolithic microfabricated valves and pumps by multilayer soft lithography. Science 288, 113–116 (2000).

## Methods References

54 Brenner, S. The genetics of Caenorhabditis elegans. Genetics 77, 71–94 (1974).

55 Frøkjær-Jensen, C. et al. Single-copy insertion of transgenes in Caenorhabditis elegans. Nature genetics 40, 1375–1383 (2008).

56 del Campo, A. & Greiner, C. SU-8: a photoresist for high-aspect-ratio and 3D submicron lithography. Journal of Micromechanics and Microengineering 17, R81 (2007).

57 Unger, M. A., Chou, H.-P., Thorsen, T., Scherer, A. & Quake, S. R. Monolithic microfabricated valves and pumps by multilayer soft lithography. Science 288, 113–116 (2000).

